# Hülle-cell-mediated protection of fungal reproductive and overwintering structures against fungivorous animals

**DOI:** 10.1101/2021.03.14.435325

**Authors:** Li Liu, Benedict Dirnberger, Oliver Valerius, Enikő Fekete-Szücs, Rebekka Harting, Christoph Sasse, Daniela E. Nordzieke, Stefanie Pöggeler, Petr Karlovsky, Jennifer Gerke, Gerhard H. Braus

**Affiliations:** University of Göttingen, Molecular Microbiology and Genetics and Göttingen Center for Molecular Biosciences (GZMB), 37077 Göttingen, Germany; University of Göttingen, Genetics of Eukaryotic Microorganisms and Göttingen Center for Molecular Biosciences (GZMB), 37077 Göttingen, Germany; University of Göttingen, Molecular Phytopathology and Mycotoxin Research, 37077 Göttingen, Germany

## Abstract

Fungal Hülle cells with nuclear storage and developmental backup functions are reminiscent of multipotent stem cells. In the soil, Hülle cells nurse the overwintering fruiting bodies of *Aspergillus nidulans*. The genome of *A. nidulans* harbors genes for the biosynthesis of xanthones. We show that enzymes and metabolites of this biosynthetic pathway accumulate in Hülle cells under the control of the regulatory velvet complex, which coordinates development and secondary metabolism. Deletion strains blocked in the conversion of anthraquinones to xanthones are delayed in maturation and growth of fruiting bodies. Xanthones are not required for sexual development but exert antifeedant effects on fungivorous animals such as springtails and woodlice. These findings reveal a novel role of Hülle cells in establishing secure niches for *A. nidulans* by accumulating metabolites with antifeedant activity that protect reproductive structures from animal predators.

## Introduction

Fungi are sessile organisms and cannot escape when they are attacked by predators or competitors. Whereas vertebrates have a protective immune system, which is regulated by Rel homology domain transcription factors, fungi have developed chemical defense strategies by producing protective secondary metabolites (SMs), which are regulated by the structurally similar velvet domain proteins (Ahmed et al., 2013). These small molecule (< 1000 Da) fungal SMs are not directly involved in the normal growth of the producing organisms but play important roles in the organism’s survivability in nature. Many fungal SMs affect the growth, survival and reproduction of surrounding organisms and are toxic or deterrent to animals (Künzler et al., 2018; Rohlfs et al., 2011). Former studies showed that the presence of bacteria or the predation by animals triggers fungal secondary metabolite production (Fischer et al., 2018; Volker Schroeckh et al., 2014; Xu et al., 2019). Secondary metabolism and development are interconnected processes (Ö. Bayram et al., 2016; Ö. Bayram et al., 2008; Ö. S. Bayram et al., 2010; Keller et al., 2019). Many fungal SMs possess intrinsic functions by being incorporated into developmental structures or functioning as signals to initiate developmental processes. For example, fungal SMs as signaling hormones induce the formation of spores and regulate their germination (Niu et al., 2020; Rodríguez-Urra et al., 2012). In addition, SMs required for the formation of fungal resting or sexual structures have been identified (Calvo et al., 2015; Schindler et al., 2014; Studt et al., 2012).

The SMs epishamixanthone and shamixanthone were commonly isolated from filamentous *Aspergillus* spp. (A. Chen et al., 2016). The cosmopolitan fungal genus *Aspergillus* comprises more than 300 species with high relevance for biotechnology, pathogenicity and post-harvest crop protection (Samson et al., 2014). The biosynthetic pathway for epishamixanthone and shamixanthone was firstly identified in *Aspergillus nidulans* (Sanchez et al., 2011) and the corresponding gene cluster consists of one polyketide synthase (PKS) encoding gene *mdpG* and 11 “tailoring” genes (*mdpA*-*F, mdpH*-*L*). The *mdp* genes are biosynthetically linked with the three *xpt* genes *xptA, xptB* and *xptC*, which are distributed over the genome, and encode two prenyltransferases and an oxidoreductase (Fig. S1). All biosynthetic genes will be referred to as the *mdp*/*xpt* gene cluster. In total, more than 30 compounds belonging mostly to the chemical groups of anthraquinones, benzophenones and xanthones are synthesized (Caesar et al., 2020; Chiang et al., 2010; Pockrandt et al., 2012; Sanchez et al., 2011). In *A. nidulans* wildtype A4, 10 out of 15 *mdp*/*xpt* genes were found up-regulated in a transcriptome analysis under sexual development inducing conditions and the three corresponding metabolites shamixanthone, emericellin and emodin were detected in a metabolome analysis from sexual cultures (Ö. Bayram et al., 2016). SM producing fungi possess global regulation mechanisms to control the SM production at specific time and in a certain tissue for particular biochemical roles (Keller et al., 2015). Whether the *mdp*/*xpt* gene cluster expression and corresponding metabolites play roles in the sexual development of *A. nidulans* remains unknown.

*A. nidulans* is a soil-borne filamentous fungus with a well-characterized life cycle (Park et al., 2019). The velvet complex VelB-VeA-LaeA accurately regulates cell differentiation and secondary metabolism in response to environmental stimuli (Ö. Bayram et al., 2008). After spore germination, a network of vegetative hyphae is formed, which develops in certain environmental conditions through asexual or sexual developmental programs to spore-bearing conidiophores or sexual fruiting bodies (cleistothecia) (Busch et al., 2007; Etxebeste et al., 2010). Light stimulates the asexual pathway, whereas lowered oxygen levels and darkness stimulate the sexual pathway, respectively (Ö. Bayram et al., 2016). The cleistothecium serves as overwintering structure and contains more than 10,000 sexual ascospores. It is surrounded by several layers of globose Hülle cells, which have nuclear storage and developmental backup functions and nurse the young fruiting body (Troppens et al., 2020). Lack of the epigenetic global regulator LaeA results in a loss of Hülle cells and in cleistothecia of reduced sizes (Ö. S. Bayram et al., 2010), whereas lack of the velvet proteins VelB and VeA abolishes cleistothecia and Hülle cells (Ö. Bayram et al., 2008; Kim et al., 2002).

We investigated the localization of the *mdp*/*xpt* cluster encoded proteins and their SM products during sexual development of *A. nidulans*. The Mdp/Xpt proteins are localized in sexual mycelia and Hülle cells, and the SMs are produced as soon as Hülle cells are present and cleistothecia begin to form. Furthermore, loss of the regulatory velvet complex proteins impaired the metabolite production of the *mdp*/*xpt* cluster. Strains with a disturbed biosynthetic pathway due to *mdp*/*xpt* gene deletions cannot produce the final epi-/shamixanthone. Instead, they accumulate various intermediates in Hülle cells, leading to smaller Hülle cells with reduced activity and a delayed maturation of cleistothecia. Therein, the accumulated intermediate emodin and its derivatives exhibit repression on sexual fruiting body and resting structure formation of other fungi. We showed in a food choice experiment that the *mdp*/*xpt* cluster metabolites present in wildtype protect *A. nidulans* from soil animal predators. These results suggest that the *mdp*/*xpt* metabolites produced in wildtype Hülle cells protect the sexual fruiting body of *A. nidulans* from fungivorous animals.

## Results

### Proteins encoded by the *mdp*/*xpt* cluster are located in Hülle cells and sexual mycelia in *Aspergillus nidulans*

Most of the *mdp*/*xpt* genes in *A. nidulans* are expressed during sexual development (Ö. Bayram et al., 2016). A comparative proteome study on protein extracts of whole sexual tissues as well as enriched Hülle cells from wildtype A4 was conducted and indicated a specific spatial and temporal accumulation of Mdp/Xpt proteins (Fig. S2a and Proteomic MS analysis data). Vegetative and asexual mycelia were used as controls. Vegetative mycelia were cultivated 20 h in liquid medium and asexual and sexual tissues as well as Hülle cells were harvested three, five and seven days after inoculation on plates. An LC-MS analysis revealed that 24 proteins were present exclusively in both sexual mycelia and enriched Hülle cells but were not identified from vegetative or asexual tissues (Table S1). Among them, five proteins encoded by the *mdp*/*xpt* cluster were identified, MdpG, MdpL, MdpH, XptB and XptC (Table 1). To verify the localization of these proteins, the final enzyme in the biosynthesis of epi-/shamixanthone, XptC, was selected as an example and C-terminally fused to GFP for fluorescence microscopy. The fusion protein XptC-GFP was exclusively detected in three days-old sexual hyphae as well as in enriched Hülle cells but not in 20 hours-old vegetative hyphae (Fig. 1a, Fig. S2b). The stability of the fusion protein XptC-GFP was verified by the α-GFP antibody in Western analysis (Fig. 1b). These results suggest that at least five members of the *mdp*/*xpt* cluster, MdpG, MdpL, MdpH, XptB and XptC, are specifically located to Hülle cells as well as sexual hyphae. Members of the Mdp/Xpt proteins can be detected from three to seven days of sexual development.

**Table 1.** Five Mdp/Xpt proteins are found in sexual mycelia and Hülle cells. *A. nidulans* wildtype A4 was sexually cultivated on MM agar plates for three, five and seven days.
Proteins were identified by LC-MS. Only proteins identified in at least two biological replicates and with two or more peptides were considered for the analysis. Numbers represent the average of spectral counts from three biological replicates. Table 1-Source data1

| Gene ID (protein) | Putative function (Szwalbe et al., 2019) | 3 d |  | 5 d |  | 7 d |  |
| --- | --- | --- | --- | --- | --- | --- | --- |
|  |  | sex. mycelia | Hülle cells | sex. mycelia | Hülle cells | sex. mycelia | Hülle cells |
| AN10023 (MdpL) | Baeyer-Villiger monooxygenase | 121 | 25 | 98 | - | 34 | 42 |
| AN7998 (XptC) | Reductase | 54 | 7 | 48 | 6 | 26 | - |
| AN12402 (XptB) | Prenyltransferase | 52 | 5 | 19 | - | - | 8 |
| AN10022 (MdpH) | Anthrone oxidase, decarboxylase | 27 | - | 2 | 5 | 27 | 6 |
| AN0150 (MdpG) | Polyketide synthase | 12 | 4 | - | 3 | - | - |

**Fig. 1.**
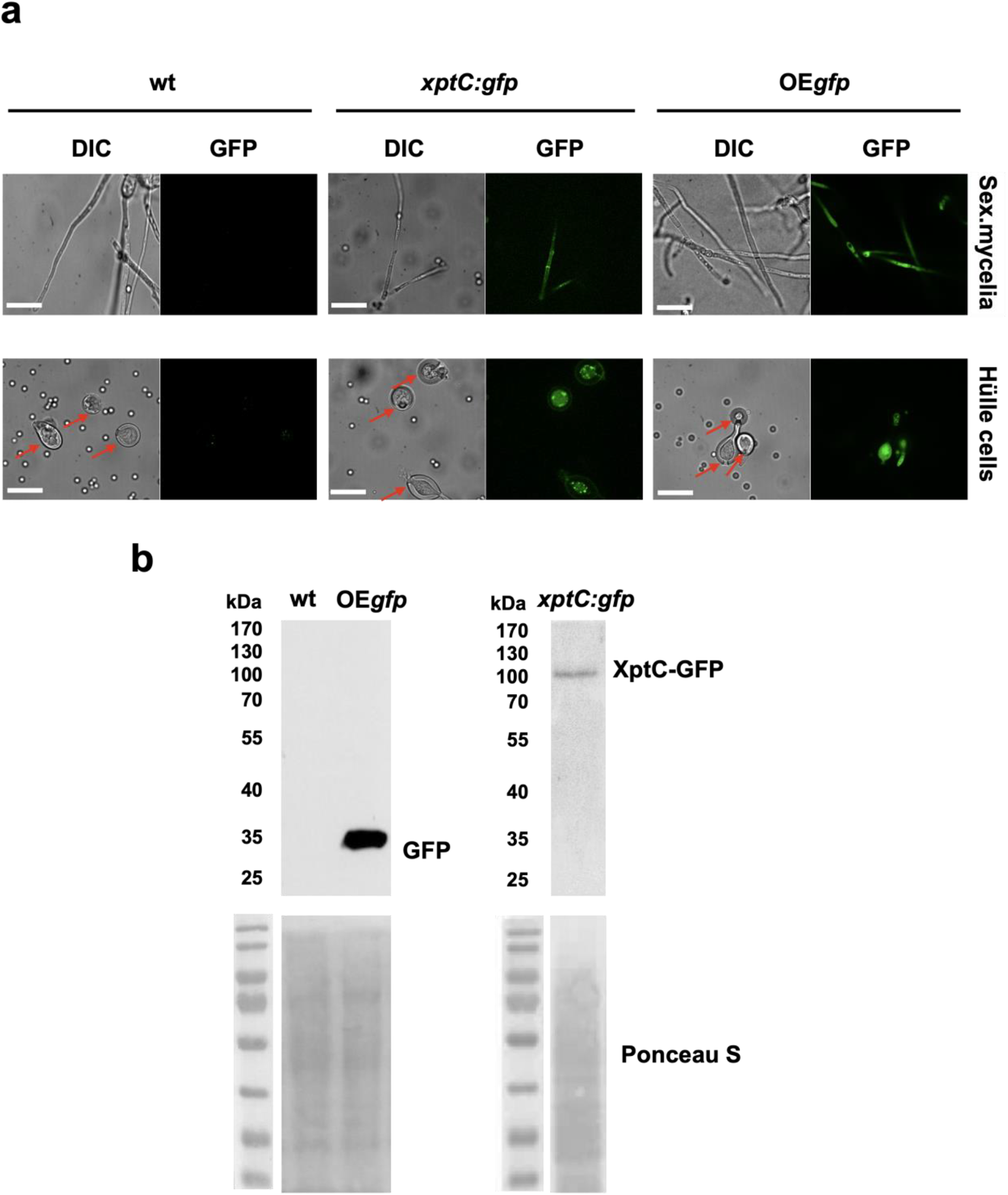
XptC is enriched in Hülle cells. **a)** Fluorescence microscopy of sexual mycelia and Hülle cells from *xptC:gfp* strain after three days of incubation. *A. nidulans* wildtype (wt, AGB552) and constitutively expressed GFP strain (OE*gfp*, AGB596) were used as controls. Red arrows indicate Hülle cells. Scale bar = 20 µm. **b**) Western hybridization of sexual (sex) tissues harvested three days after inoculation. α-GFP antibody was used. The fusion protein XptC-GFP was detected at approximately 95 kDa. A strain expressing GFP constitutively (OE*gfp*, AGB596) and wildtype (wt, AGB552) served as controls. Ponceau S staining was used as sample loading control.

### The concentration of *mdp*/*xpt* cluster metabolites changes over time with fruiting body development

Under laboratory conditions, *A. nidulans* wildtype (AGB552), harboring Δ*nkuA* for improved homologous gene replacements, forms young sexual fruiting bodies (cleistothecia) that are surrounded by Hülle cells after three days of sexual growth in the absence of light. At this stage, all *mdp*/*xpt* genes are expressed (Fig. S3). After five days, the cleistothecia are mature with a dark pigmented shell (Fig. S4). Timing and localization of *mdp*/*xpt* metabolite production were monitored during sexual development and their roles in cleistothecia formation were examined. Different time points were selected for SM analysis (Fig. S4): prior to the development of cleistothecia or Hülle cells (day 2), young cleistothecia with Hülle cells (day 3) and mature cleistothecia with Hülle cells (day 5), as well as late stages with mature cleistothecia (day 7 and 10). The different SM intermediates were addressed by a genetic approach. The pathway was disturbed by deleting *mdpG* and *mdpF* (early biosynthetic steps), *mdpC* and *mdpL* (intermediate biosynthetic steps) and *mdpD, xptA, xptB* and *xptC* (late biosynthetic steps) separately (Fig. 2A). *A. nidulans* wildtype and *mdp*/*xpt* deletion strains were grown under sexual conditions for two, three, five, seven and 10 days and the extra- and intracellular metabolites were extracted with ethyl acetate and subjected to LC-MS analysis. The identified *mdp*/*xpt* cluster products were categorized into four groups according to their molecular structures (anthraquinones in orange, benzophenones in blue, xanthones in green, arugosins in black, Fig. 2a).

**Fig. 2.**
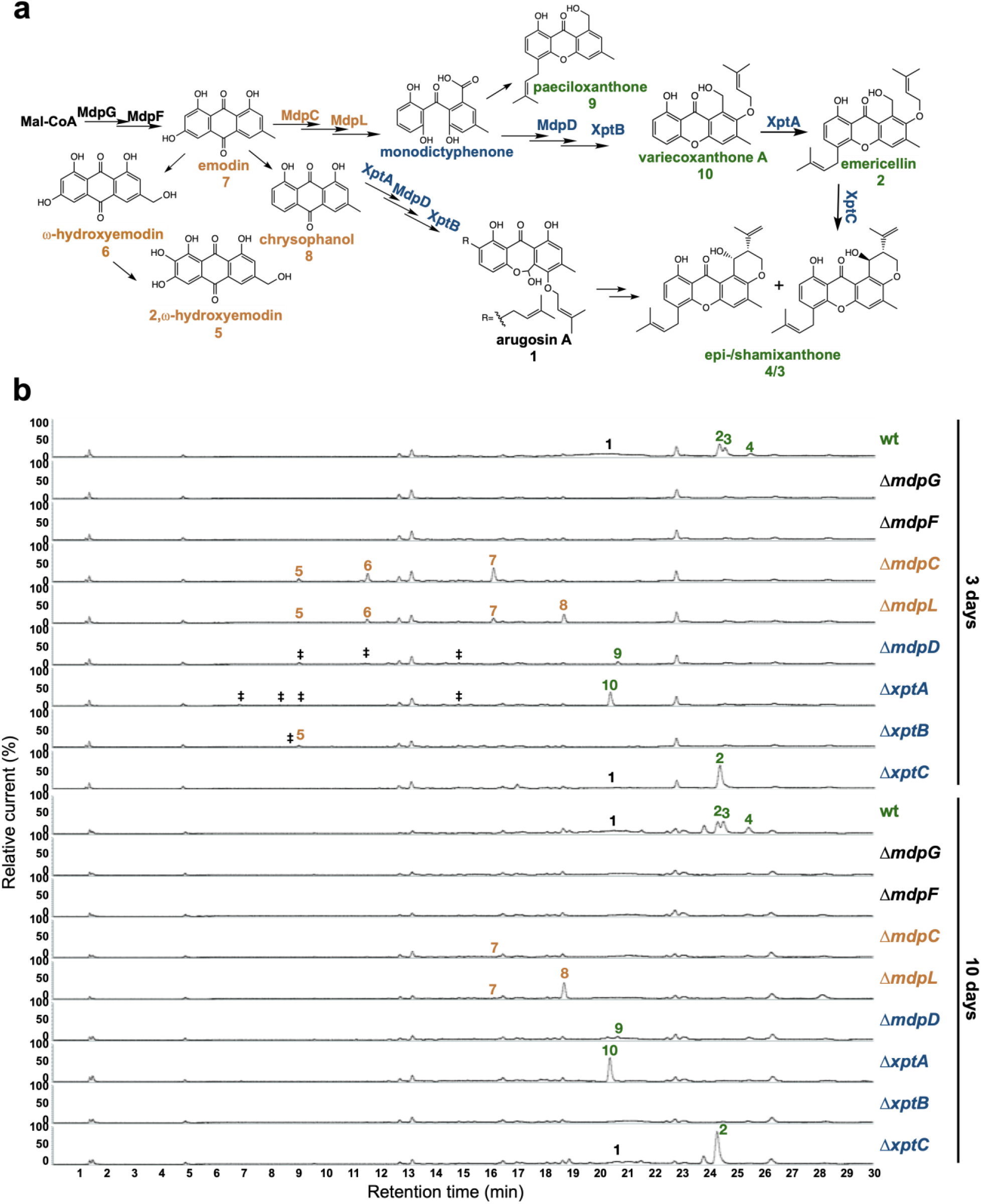
The *mdp*/*xpt* cluster metabolites are produced after three days of sexual growth in *A. nidulans*. **a**) Simplified biosynthetic pathway of epi-/shamixanthone in *A. nidulans* (Chiang et al., 2010; Pockrandt et al., 2012; Sanchez et al., 2011). Black enzymes are localized at the early steps of the biosynthetic pathway. Orange enzymes are localized at the middle of the biosynthetic pathway. Blue enzymes are localized at the late steps of the biosynthetic pathway. The cluster products are classified as four groups: anthraquinones are orange, benzophenones are blue, xanthones are green, arugosin A is black. **b**) Chromatograms of secondary metabolites (SMs) of *A. nidulans* wildtype (wt, AGB552) and *mdp*/*xpt* mutant strains (Δ). Conidia of *A. nidulans* wt and *mdp*/*xpt* mutant strains were point-inoculated on minimal medium (MM) and sexually grown for three and 10 days. Extra- and intracellular metabolites were extracted and detected by LC-MS with a charged aerosol detector (CAD) in three independent experiments. Only CAD-detectable and identified SMs are shown with numbers. **1**) arugosin A; **2**) emericellin; **3**) shamixanthone; **4**) epishamixanthone; **5**) 2,ω-dihydroxyemodin; **6**) ω-hydroxyemodin; **7**) emodin; **8**) chrysophanol; **9**) paeciloxanthone; **10**) variecoxanthone A. **‡** marks unidentified compounds.

After two days of sexual development, wildtype and deletion strains did not produce any *mdp*/*xpt* metabolites (Fig. S5). After three and five days, wildtype produced arugosin A (**1**) and the final xanthones emericellin (**2**), shamixanthone (**3**) and epishamixanthone (**4**) (Fig. 2b, Fig. S5, Table S2). As expected, loss of the first two enzymes of the biosynthesis MdpG and MdpF completely abolished the production of cluster metabolites. Deletion of the intermediate enzyme encoding genes *mdpC* and *mdpL* led to a loss of **1**-**4** but to the accumulation of the anthraquinones 2,ω-dihydroxyemodin (**5**), ω-hydroxyemodin (**6**), emodin (**7**) in both strains and chrysophanol (**8**) in Δ*mdpL*. Deleting the biosynthetically final enzyme encoding genes *mdpD, xptA* and *xptB* abolished the production of **1**-**4**. Δ*mdpD* and Δ*xptA* accumulate the xanthones paeciloxanthone (**9**) and variecoxanthone A (**10**), respectively. Δ*xptB* accumulates the anthraquinone **5**. In addition, these three strains accumulate some unidentified compounds. Deletion of *xptC* led to a loss of the final xanthones **3** and **4** but an increased accumulation of **2**. After seven and 10 days of sexual development, the accumulated emodins **5-7** of the deletion strains were decreased or even disappeared (Fig. 2b, Fig. S5), whereas the abundance of **8** increased and the xanthones **2-4** and **10** were still detectable in similar amounts as after 3 and 5 days.

In summary, the SMs of the *mdp*/*xpt* cluster are produced as soon as the first sexual structures are visible after three days of sexual development. During cleistothecia maturation from three to 10 days, the amount of emodins gradually decreased, whereas the amount of chrysophanol and the xanthones increased or remained stable.

### The *mdp*/*xpt* cluster metabolites are enriched in Hülle cells of *A. nidulans*

The colors of the *mdp*/*xpt* cluster metabolites mostly are yellow or orange in pure powder form (Chiang et al., 2010). This assisted us to trace the metabolites localization in the fungus. *A. nidulans* wildtype forms a colony with green conidiospores and light yellow Hülle cells after three days of sexual growth (Fig. 3a and b). All *mdp*/*xpt* deletion strains showed no change in green conidiospore production but the color of the Hülle cells changed. Whereas Δ*mdpD*, Δ*xptA*, Δ*xptB* and Δ*xptC* showed no color difference to wildtype, the deletion strains of the early biosynthetic genes *mdpG* and *mdpF*, which have lost *mdp*/*xpt* metabolite synthesis (Fig. 2b), produced colorless Hülle cells. Δ*mdpC*, which accumulates the brown (**5**) and yellow (**6, 7**) emodins and yellow chrysophanol (**8**), produced dark yellow Hülle cells (Fig. 3a and b). This indicates that the *mdp*/*xpt* metabolites are enriched inside the Hülle cells. The SMs have no obvious effect on Hülle cell shape (Fig. 3c) but the accumulation of epi-/shamixanthone precursors in Δ*mdpC*, Δ*mdpL*, Δ*mdpD*, Δ*xptA*, and Δ*xptB* decreased the Hülle cell size (Fig. 3d). The strains with the smallest Hülle cells, Δ*mdpC* and Δ*mdpL*, were tested for their Hülle cell germination ability. 39-56% of tested Hülle cells of wildtype and Δ*mdpG* germinated, whereas the small Hülle cells of Δ*mdpC* and Δ*mdpL* displayed a germination ability of only 2-6% (Fig. 3e).

**Fig. 3.**
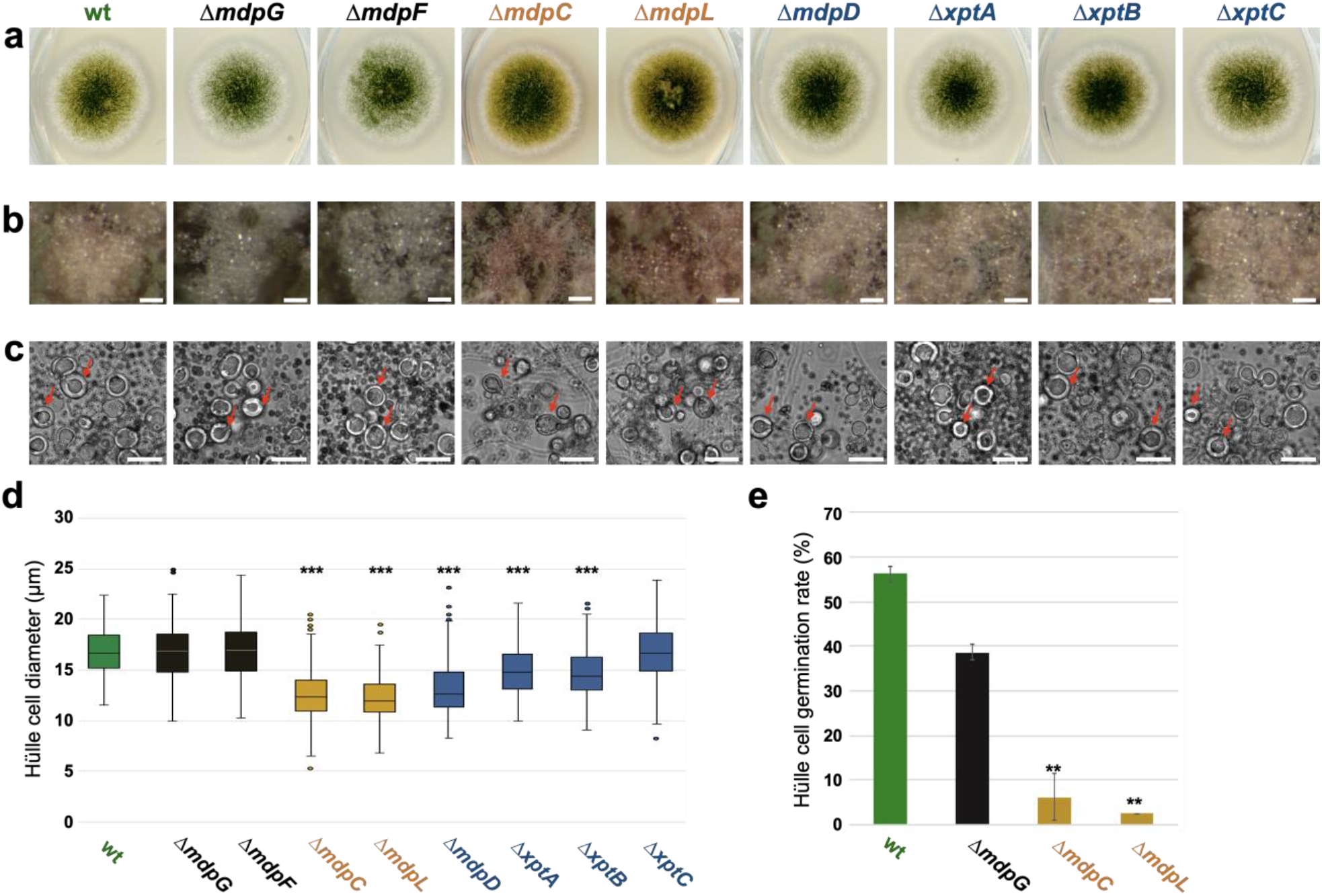
The *mdp*/*xpt* cluster metabolites are localized in Hülle cells. **a**) Colony phenotypes of *A. nidulans* wildtype (wt, AGB552) and *mdp*/*xpt* deletion strains (Δ). Conidia were point-inoculated on MM agar plates and cultivated three days under sexual conditions. **b**) Photomicrographs of Hülle cells after five days. Scale bar = 50 µm. **c**) Morphology of Hülle cells after five days. Red arrows indicate examples of Hülle cells. Scale bar = 25 µm. **d**) Box plot of Hülle cells size after five days of sexual development (n ≥ 150). **e**) Germination rate of Hülle cells. Detached Hülle cells were collected from cleistothecia surface after five days of sexual development and placed on fresh MM agar plates. The germination was monitored after 48 hours at 37°C. n = 40 (± 1) with two biological replicates. All significance tests are in comparison to wildtype (wt), ***/ ** *P* < 0.005 / 0.05, two-tailed *t* test. Figure 3d-Source data 2 Figure 3e-Source data 3

### The velvet complex regulates the *mdp*/*xpt* cluster metabolite production in *A. nidulans*

Velvet proteins, such as VeA and VelB, are fungal DNA-binding proteins with a similar structural fold as the mammalian NF-κB inflammation and infection regulators (Ahmed et al., 2013). In *A. nidulans*, velvet proteins physically interact with epigenetic methyltransferases like LaeA (Sarikaya-Bayram et al., 2014; Sarikaya-Bayram et al., 2015). The heterotrimeric velvet complex VelB-VeA-LaeA coordinates sexual development and secondary metbolism (Ö. Bayram et al., 2008). VelB-VeA enters the nucleus to initiate sexual development and physically interacts with the epigenetic master regulator of secondary metabolism LaeA. In order to analyze the impact of the velvet complex on *mdp*/*xpt* SM production, extra- and intracellular metabolites of wildtype, Δ*veA*, Δ*velB* and Δ*laeA* were analyzed by LC-MS after five days of sexual development (Fig. 4). The production of the final *mdp*/*xpt* products, arugosin A (**1**) and the xanthones emericellin (**2**), shamixanthone (**3**) and epishamixanthone (**4**), was abolished in Δ*veA* and Δ*velB* and was reduced in Δ*laeA*. This indicates that the velvet complex VelB-VeA-LaeA regulates the cluster metabolite production, whereby VelB and VeA are prerequisites for the *mdp*/*xpt* metabolite production.

**Fig. 4.**
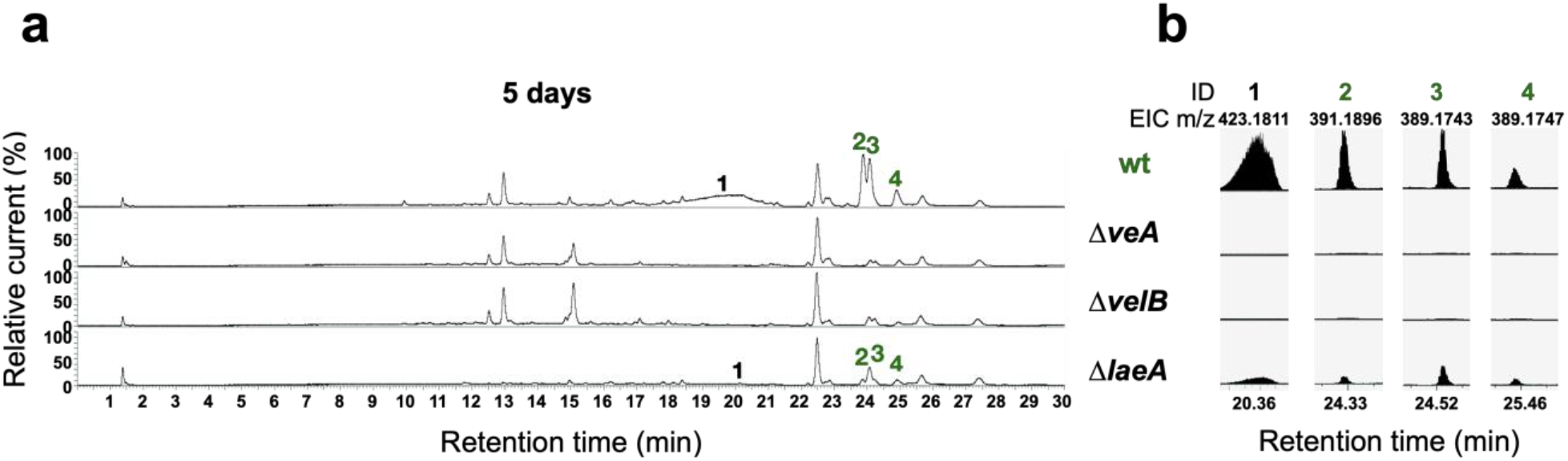
The velvet complex is required to produce *mdp*/*xpt* cluster metabolites. **a)** Chromatograms of secondary metabolites (SMs) of *A. nidulans* wildtype (wt, AGB552) and velvet complex gene deletion strains (Δ). Conidia of *A. nidulans* strains were point-inoculated on minimal medium (MM) and sexually grown for five days. Extra- and intracellular metabolites were extracted and detected by LC-MS with a charged aerosol detector (CAD). Only CAD-detectable SMs of the *mdp*/*xpt* cluster are shown with numbers and were identified with MS and UV/VIS. **b**) EICs (extracted ion chromatograms) of the compounds detected by CAD. m/z of **1** was detected in negative mode. m/z of **2, 3** and **4** was detected in positive mode. ID (compound number in this study): **1**) arugosin A; **2**) emericellin; **3**) shamixanthone; **4**) epishamixanthone.

### The intact *mdp*/*xpt* cluster is required for accurate sexual development in *A. nidulans*

Wildtype of *A. nidulans* forms young, unpigmented cleistothecia after three days that are covered with high amounts of Hülle cells. Cleistothecia shells become pigmented after four days, and cleistothecia are mature with a dark pigmented shell after five days (Fig. 5a). Development of sexual structures was monitored over time in wildtype and *mdp*/*xpt* deletion strains in order to get insights into the biological roles of the *mdp*/*xpt* cluster metabolites. Deletion strains of *mdpG* and *mdpF*, which produce no *mdp*/*xpt* SMs (Fig. 2b), developed wildtype-like cleistothecia, indicating that the final *mdp*/*xpt* xanthones and arugosin A are not required for sexual development. In contrast, Δ*mdpC* and Δ*mdpL*, which both accumulate the emodins **5**-**7**, are strongly delayed in cleistothecia development. After three days of sexual development, they formed pigmented Hülle cells without any cleistothecia. After five days, young, immature cleistothecia with soft and barely pigmented shells were formed. Cleistothecia were fully pigmented after 10 days (Fig. 5a) with reduced sizes (Fig. 5b) and diminished numbers of ascospores (Fig. 5c), but with viable ascospores (Fig. S6). Even after 25 days, cleistothecia size was still significantly smaller than for wildtype (Fig. S7). For Δ*mdpL*, the number of cleistothecia was significantly increased after 10 days compared to all other strains (Fig. 5d). The deletion strains of *mdpD, xptA* and *xptB* exhibited a moderate delay of cleistothecia development (Fig. 5a). After four days of sexual development, they formed young, only barely pigmented cleistothecia, but after five days the delay was rescued and the cleistothecia were wildtype-like. In addition, for Δ*xptB*, cleistothecia diameter (Fig. 5b and Fig. S7) as well as ascospore amount (Fig. 5c) were diminished, but the spores were viable (Fig. S6).

**Fig. 5.**
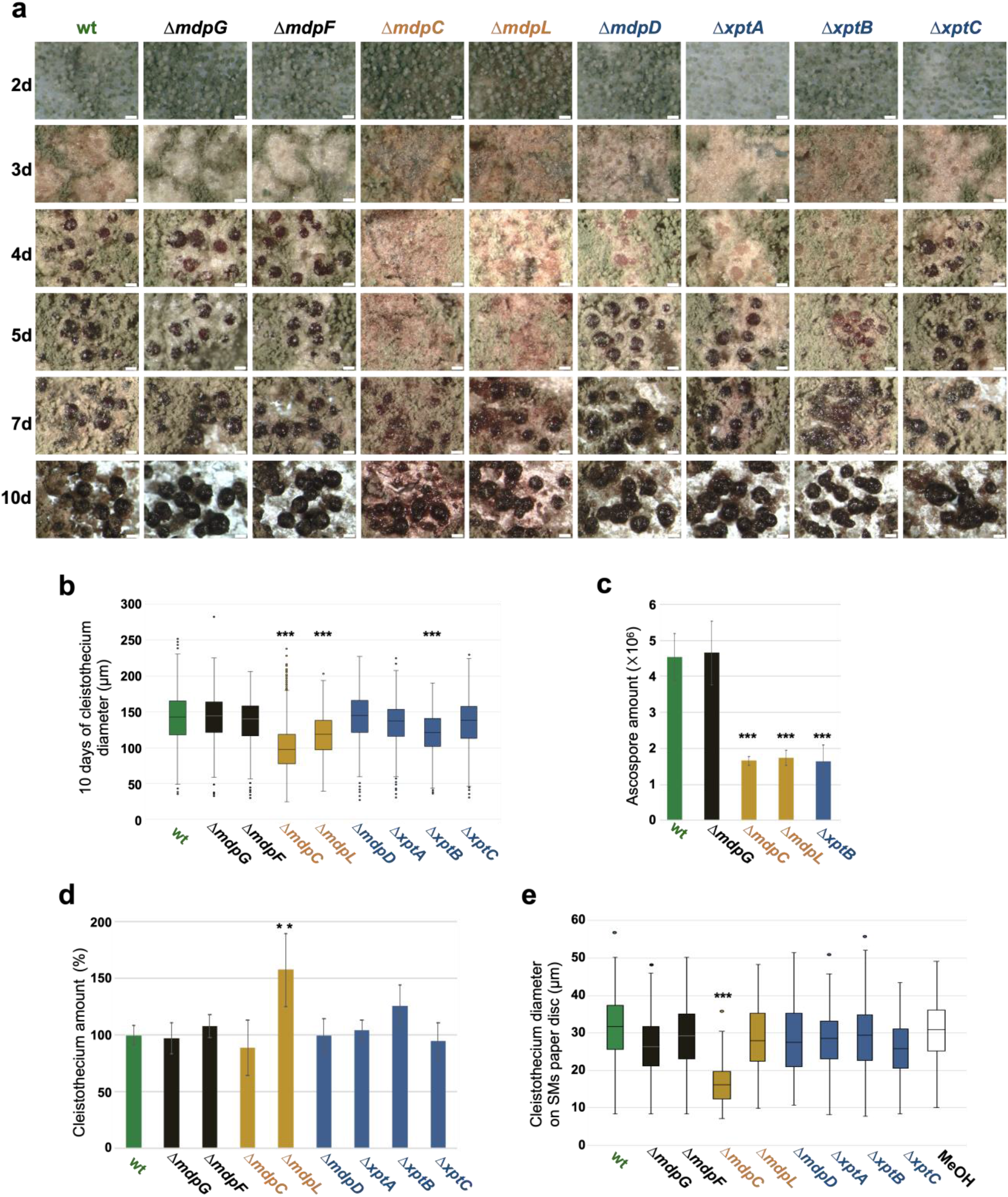
Accumulated intermediates of the *mdp*/*xpt* cluster repress sexual development of *A. nidulans*. **a**) Photomicrographs of sexual structures of wildtype (wt, AGB552) and *mdp*/*xpt* deletion strains at different developmental stages. Scale bar = 100 µm. **b**) Box plot of cleistothecia diameter after 10 days of sexual development (n ≥ 650). **c**) Ascospore quantification after 10 days. 10 cleistothecia were broken in 100 µl 0.02% tween buffer and released ascospores were quantified. Error bar indicates standard deviation with three biological and three technical replicates. **d**) Cleistothecia quantification after 10 days. Error bar indicates standard deviation with five biological replicates; amount of cleistothecia of wt was set to 100%. **e**) Box plot of cleistothecia diameter of *A. nidulans* wt grown on paper discs loaded with extracted SMs of wt and *mdp*/*xpt* deletion strains. Strains were sexually grown for five days. SMs were extracted, solved in MeOH and loaded on paper discs separately (pure MeOH was used as blank control). Paper discs were placed on agar plates inoculated with 1 × 10^5^ conidia of *A. nidulans* wt. Cleistothecia on the paper discs were collected after five days of sexual growth (n ≥ 65). All significance tests are in comparison to wt, ***/ ** *P* < 0.005 / 0.05, two-tailed *t* test. Figure 5b-Source data 4 Figure 5c-Source data 5 Figure 5d-Source data 6 Figure 5e-Source data 7

The effect of the extracted metabolites of the *mdp*/*xpt* deletion strains on cleistothecia development of *A. nidulans* wildtype was examined to analyze whether the delayed maturation of cleistothecia correlates with the accumulated cluster metabolites. Metabolites of wildtype and *mdp*/*xpt* deletion strains were extracted after five days of sexual development, solved in methanol and loaded onto paper discs separately. SM loaded paper discs were placed on an agar plate inoculated with *A. nidulans* wildtype spores. Pure methanol served as control. After five days of sexual incubation, the cleistothecia development on each paper disc was monitored. Metabolites of wildtype and Δ*mdpG*, Δ*mdpF*, Δ*mdpL*, Δ*mdpD*, Δ*xptA*, Δ*xptB* and Δ*xptC* displayed no effect on cleistothecia development. Metabolites of Δ*mdpC* exhibited obvious effects on the wildtype cleistothecia development producing significantly smaller and immature cleistothecia (Fig. 5e and Fig. S8). This verified that the metabolites of Δ*mdpC* 2,ω-dihydroxyemodin (**5**), ω-hydroxyemodin (**6**) and emodin (**7**), or at least one of them, negatively influence cleistothecia development. In summary, the incomplete epi-/shamixanthone biosynthetic pathway causes the accumulation of intermediates, in particular of emodins, resulting in a delay in cleistothecia maturation and a repression of cleistothecia size. This indicates that an intact *mdp*/*xpt* pathway is required for accurate sexual development in *A. nidulans* and an interrupted xanthone biosynthesis negatively influences cleistothecia development.

### Epi-/shamixanthone precursors in *A. nidulans* repress fruiting body and resting structure formation of other fungi

The soil fungus *A. nidulans* forms cleistothecia as overwintering structures to survive in harsh environments until its favorable growth conditions return. Enriched epi-/shamixanthone precursors in *mdp*/*xpt* deletion strains showed a repression on cleistothecia development, which might give other soil fungi a competitive advantage for their own fruiting body or resting structure formation. We analyzed the effect of those precursors on sexual reproduction or resting structures formation of other fungi to examine whether *A. nidulans* can make use of these chemicals to compete against other fungi for its further development. The effects on sexual reproduction of the fungus *Sordaria macrospora* was tested, which lacks an asexual reproduction cycle. Further, the effects of epi-/shamixanthone precursors on melanized resting structures of two asexually reproducing *Verticillium* spp were investigated. SMs extracted from *A. nidulans* wildtype and *mdp*/*xpt* deletion strains were loaded for all approaches onto paper discs and placed on agar plates inoculated with spores of the corresponding fungus.

*S. macrospora* produces flask-shaped, pigmented sexual fruiting bodies (perithecia) after seven days of surface cultivation (Teichert et al., 2020). When *S. macrospora* was exposed to SMs of *A. nidulans* Δ*mdpC*, Δ*mdpL* and Δ*mdpD*, perithecia formation was repressed, resulting in a halo surrounding the SM loaded paper disc. The biggest halo was observed for Δ*mdpC* metabolites (Fig. 6, Fig. S9). *Verticillium dahliae* and *V. longisporum* form melanized hyphal aggregates called microsclerotia as resting structures to survive in the soil for decades (Zeise et al., 2001). Exposed to the SMs of *A. nidulans* Δ*mdpC*, Δ*mdpL* and Δ*mdpD, V. dahliae* and *V. longisporum* produced fewer microsclerotia under the paper discs and its surrounding area. Especially under the paper disc with SMs of Δ*mdpC, Verticillium* spp. cannot produce any microsclerotia (Fig. 6). This shows that epi-/shamixanthone precursors of Δ*mdpC*, Δ*mdpL* and Δ*mdpD* repress sexual reproduction and resting structure formation of different fungi. The strongest effect was observed for Δ*mdpC*, which produces the emodins **5, 6** and **7**.

**Fig. 6.**
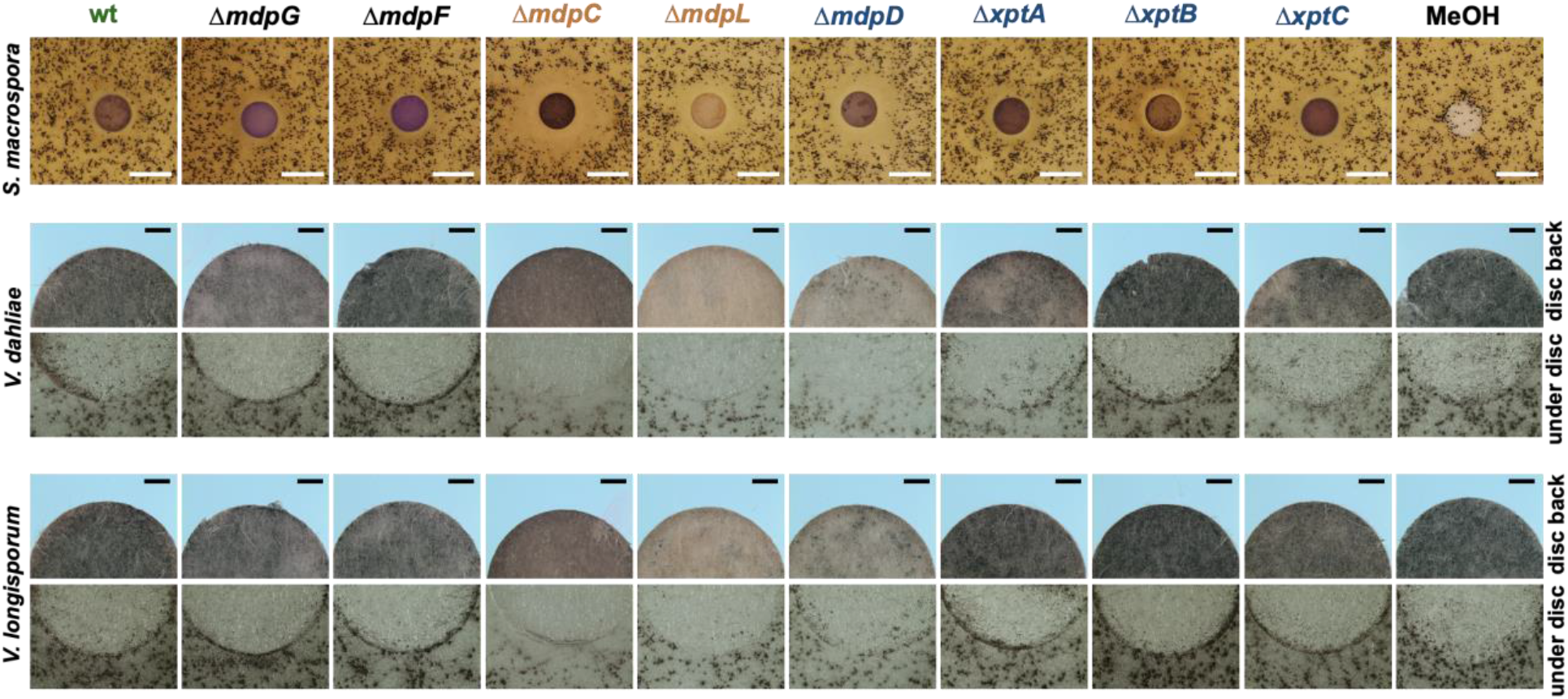
Intermediates of the *mdp*/*xpt* cluster repress fungal reproduction and resting structure formation. Microphotographs of fungal reproduction and resting structures exposed to extracted SMs of *A. nidulans* wildtype (wt) and *mdp*/*xpt* deletion strains. Conidia of *A. nidulans* strains were point-inoculated on MM and sexually grown for five days. SMs were extracted, solved in MeOH and loaded onto paper discs (MeOH solvent served as control). Paper discs were placed on agar plates inoculated with spores of the tested fungi. For *S. macrospora*, BMM agar plates were inoculated with 2 × 10^5^ spores and cultivated seven days at 27°C. White bar = 1 cm. For *Verticillium* spp., simulated xylem medium (SXM) agar plates were inoculated with 1 × 10^5^ *V. dahliae* or *V. longisporum* spores and cultivated 10 days at 25°C. The upper panel shows pictures taken from the back of the paper disc and the lower panel shows the agar under the paper disc. Black bar = 1 mm.

In order to identify the active components, commercially available pure chemicals were tested. 75 µg of ω-hydroxyemodin (**6**), emodin (**7**) and chrysophanol (**8**) were loaded separately onto paper discs and placed on agar plates inoculated with *A. nidulans, S. macrospora, V. dahliae* and *V. longisporum* wildtypes (Fig. 7). *S. macrospora, V. dahliae* and *V. longisporum* produced fewer fruiting bodies or resting structures, respectively, when exposed to emodin (**7**) in comparison to the MeOH control, ω-hydroxyemodin (**6**) and chrysophanol (**8**). The pure chemicals have no obvious effect on sexual fruiting body of *A. nidulans*. This shows that the precursor emodin, derived from the *mdp*/*xpt* cluster, suppresses fruiting body and resting structure formation of other fungi, suggesting a competitive role in nature between *A. nidulans* and other fungi.

**Fig. 7.**
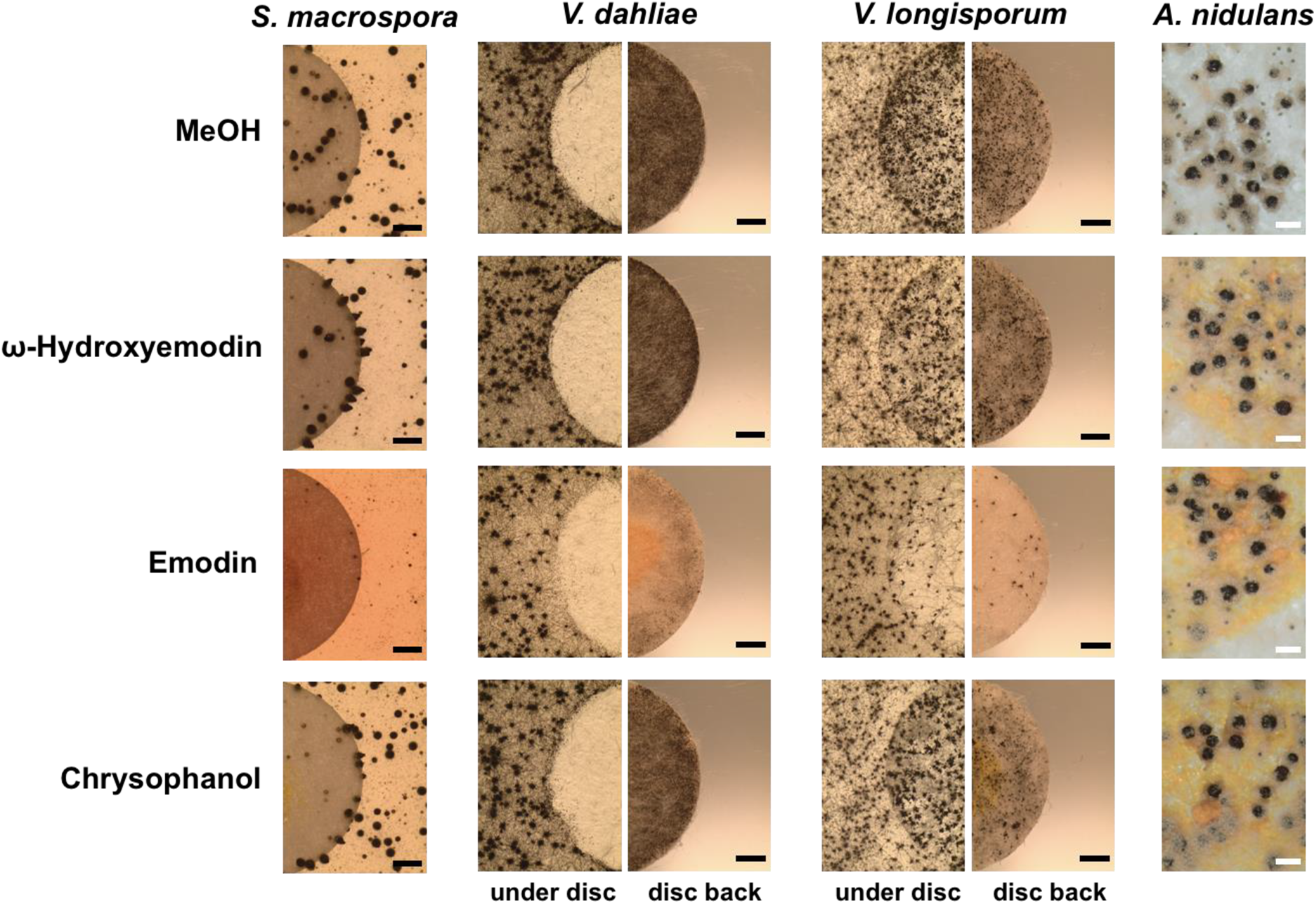
Emodin represses fungal reproduction and resting structure formation. Pure emodin, ω-hydroxyemodin and chrysophanol were dissolved in MeOH and loaded onto paper discs (final amount 75 µg per disc). MeOH served as blank control. Paper discs were placed on agar plates inoculated with spores of the tested fungi. For *S. macrospora*, BMM agar plates were inoculated with 2 × 10^5^ spores and cultivated seven days at 27°C. For *Verticillium* spp., SXM agar plates were inoculated with 1 × 10^5^ spores and cultivated 10 days at 25°C. Microsclerotia were monitored under the paper disc and at its back. Black bar = 1 mm. For *A. nidulans*, 1 × 10^5^ conidia of wildtype were inoculated on MM agar plates and sexually incubated for five days at 37°C. White bar = 200 µm.

### The final metabolite products of the *mdp*/*xpt* cluster in *A. nidulans* wildtype repel animal predators

Besides competition with other soil-borne microorganisms, fungi are facing the risk to be attacked by fungivores. Therefore, we examined whether metabolites from the *mdp*/*xpt* cluster protect *A. nidulans* from predators. The *mdpC* and *mdpG* complementation strains (*mdpC* ^*com*^, *mdpG* ^*com*^), which produce all *mdp*/*xpt* cluster metabolites like wildtype (Fig. S10), the non-producing strain Δ*mdpG* as well as the emodins accumulating strain Δ*mdpC* were offered to animal predators in a food choice experiment (Fig. 8). Animals representing distant arthropod lineages were selected: the mealworm larvae *Tenebrio molitor* (insect), the collembolan *Folsomia candida* (primitive arthropod) and the woodlouse *Trichorhina tomentosa* (crustacean). Two different fungal cultures were placed onto opposite sides of a Petri dish, and active animals were placed onto the center area. The animals feeding on each fungal culture were counted along with time.

**Fig. 8.**
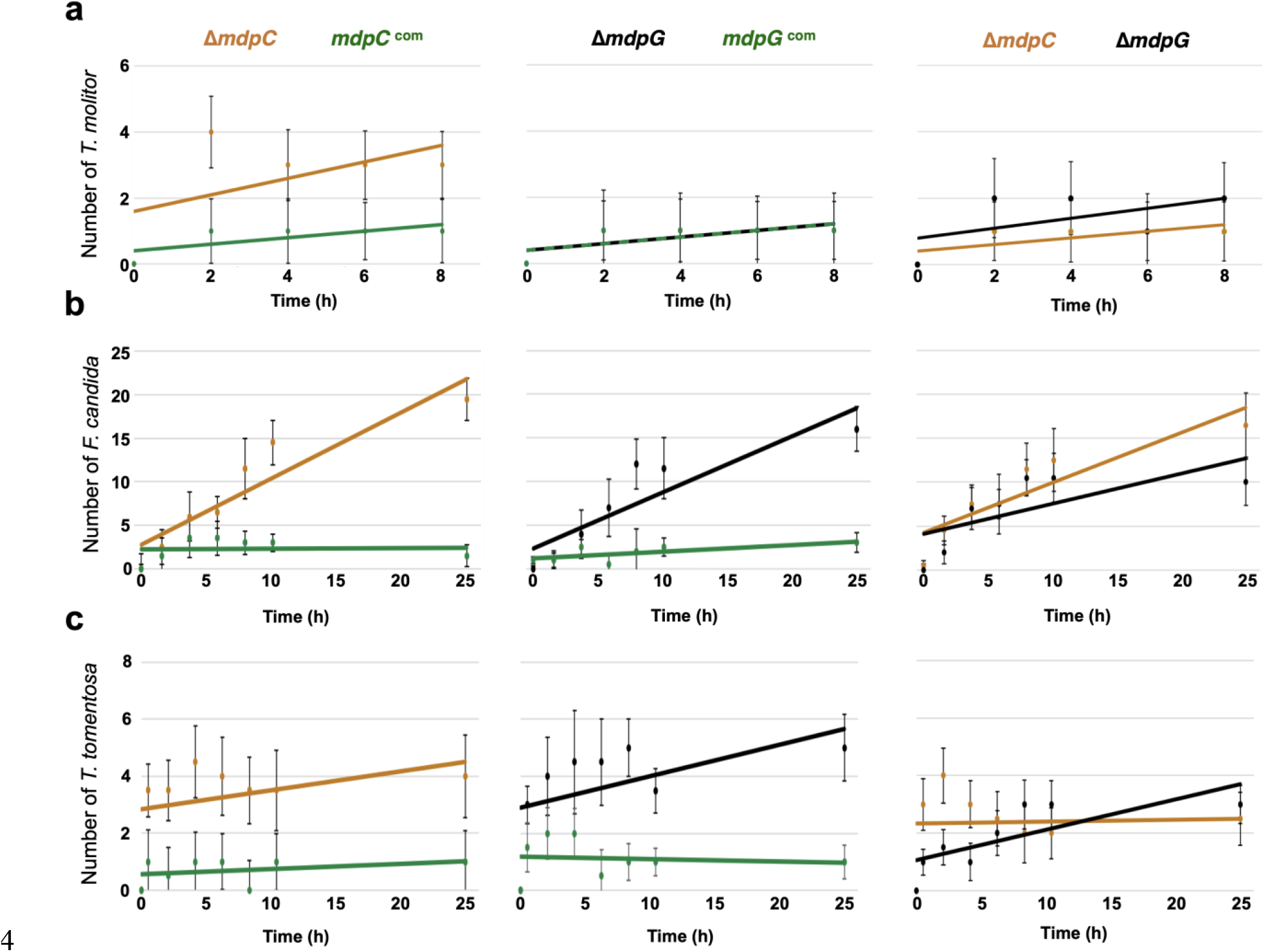
The final SM products of the *mdp*/*xpt* cluster repel fungal predators. Food preference test for *Tenebrio molitor* larvae (**a**), *Folsomia candida* (**b**) and *Trichorhina tomentosa* (**c**). *A. nidulans* spores were point-inoculated on MM agar plates and incubated for five days under sexual conditions. A piece of the colony was cut out and used for animal feeding. Animals were placed into the center of the Petri dish containing fungal agar pieces on opposite sides. The number of animals on each side was counted. For *T. molitor*, 10 animals per plate were used. The experiment was performed with three biological and nine technical replicates and medians were calculated (the medians for the number of animals of Δ*mdpG* and *mdpG* ^com^ were identical). For *F. candida*, approximately 20 animals per plate were used. For *T. tomentosa*, eight animals per plate were used. These experiments were performed with three biological and four technical replicates. Error bars represent 95% confidence interval; orange = Δ*mdpC*; black = Δ*mdpG*; green = complementation strains. Figure 8-Source data 8

*T. molitor* showed no obvious food preference facing the *mdp*/*xpt* cluster metabolites (Fig. 8a). *F. candida* and *T. tomentosa* displayed a strong avoidance for *A. nidulans* producing the final cluster products including epi-/shamixanthone (Fig. 8b-c). The springtails and isopods gradually gathered on fungal cultures not producing the final products, where they remained over 24 hours, indicating that the final cluster products deter animal predators from feeding. Emodins, which are accumulated in Δ*mdpC*, have no effect on animal grazing. A similar number of animals were found on Δ*mdpC* as on Δ*mdpG* cultures. Taken together, the production of the final *mdp*/*xpt* cluster products in *A. nidulans* have a protective function and repel animal predators such as *F. candida* and *T. tomentosa* from grazing.

## Discussion

Secondary metabolites (SMs) are small molecular products with high structural diversity enabling different biological functions. Although not directly involved in growth, SMs possess protective properties for the producing organism and are synthesized as cellular response to abiotic and biotic stimuli. They protect against environmental stresses such as oxidation and UV radiation (Narsing Rao et al., 2017) and inhibit bacterial or fungal growth (Künzler et al., 2018; Wheatley et al., 2002). Besides, SMs have antifeedant activities. Aurofusarin protects *Fusarium* species from animal grazing and asparasone A protects *Aspergillus flavus* from insect predators (Cary et al., 2014; Xu et al., 2019). The main finding of this study is that the soil fungus *A. nidulans* accumulates various xanthones in a specific fungal sexual cell type, the Hülle cells (Ö. S. Bayram et al., 2010). These xanthones are not required for sexual development but protect by exerting antifeedant effects on fungivorous animals such as springtails or woodlice.

Fruiting bodies (cleistothecia) of the soil fungus *A. nidulans* are produced as reproductive and overwintering structures and their formation is highly material- and energy-consuming (Pöggeler et al., 2006). Thus, a potent protection mechanism against predators is needed to ensure long-term survival. The animal predator *F. candida* prefers vegetative hyphae and conidia rather than cleistothecia when feeding on *A. nidulans* (Döll et al., 2013). The cleistothecia are covered with globose Hülle cells which are described to be nursing cells (Ö. S. Bayram et al., 2010) and fungal backup stem cells with nuclear storage function to produce genetically diverse spores in changing nutrient conditions (Troppens et al., 2020). Here, we showed that the *mdp*/*xpt* metabolites are produced in Hülle cells (Fig. 3) and that they deter the animal predators *F. candida* and *T. tomentosa* from feeding (Fig. 8b-c). This suggests a novel additional role for Hülle cells to preserve survival of cleistothecia. The *mdp*/*xpt* metabolites are produced as soon as the first Hülle cells occur to protect the developing, young fruiting body and are still present when the cleistothecia are mature, guaranteeing long-term protection (Fig. 2b, Fig. S5 and Fig. 5a). Fruit fly larvae prefer grazing on strains deleted for the regulatory *veA* or *laeA* genes (Trienens et al., 2010; Trienens et al., 2012). Both encoded proteins are components of the trimeric velvet complex, consisting of the two velvet-domain proteins VeA and VelB as well as the epigenetic methyltransferase LaeA.

The complex is involved in the coordinated regulation of secondary metabolism and sexual development. Deletion of either gene results in defective Hülle cell and fruiting body development and disturbed SM production (Ö. Bayram et al., 2008; Ö. S. Bayram et al., 2010). Transcriptome analysis revealed that VeA positively regulates the expression of eight out of 15 *mdp*/*xpt* genes (Lind et al., 2015). Here, deletion of the genes for VeA and VelB led to the abolishment of the *mdp*/*xpt* cluster metabolites (Fig. 4). Deletion of *laeA* reduced not only the number of Hülle cells, but also the production of cluster metabolites. These findings indicate that the production of the antifeedant *mdp*/*xpt* metabolites is under the regulatory control of the velvet complex.

The *mdp*/*xpt* gene cluster is moderately conserved in two closely related *Aspergillus* species, *Aspergillus versicolor* and *Aspergillus sydowii* (de Vries et al., 2017). *A. nidulans* and *A. versicolor* form cleistothecia, whereas a sexual cycle has not been found yet for *A. sydowii* (Raper et al., 1965), but all of them form Hülle cells (Dyer et al., 2012). The velvet complex VelB-VeA-LaeA is conserved in the fungal kingdom (Ö. Bayram et al., 2012c) and required for Hülle cell formation (Ö. S. Bayram et al., 2010; Kim et al., 2002). The strains Δ*veA* and Δ*velB* cannot produce Hülle cells and Δ*laeA* just produces a few Hülle cells. This is well connected with the *mdp*/*xpt* metabolites production in velvet complex gene deletion strains (Fig. 4) and suggests that the velvet complex regulation of the *mdp*/*xpt* cluster is Hülle cell dependent. The *Aspergillus* section *Nidulantes* is subdivided in seven clades and 65 species. Three clades and 34 species have a sexual state forming the fruiting body cleistothecium, typically surrounded by layers of Hülle cells. Therein, 27 species have been identified to produce *mdp*/*xpt* metabolites, *e*.*g*. emericellin, 2,ω-hydroxyemodin or shamixanthone (A. Chen et al., 2016). This suggests that the *mdp*/*xpt* metabolites are commonly Hülle cell-associated and widely play protective roles for the sexual reproduction of species of the genus *Aspergillus*.

The *mdp* gene cluster is biosynthetically connected with three *xpt* genes, which are distributed over the genome. The *mdp* genes produce emodin as well as monodictyphenone, whereas the *xpt* genes use the *mdp* metabolites as precursors to convert them to the final xanthones and arugosin A (Fig. 2a) (Chiang et al., 2010; Pockrandt et al., 2012; Sanchez et al., 2011). We found biological functions not only for the final *mdp*/*xpt* products but also for the *mdp* metabolite emodin. Emodin represses the formation of fruiting bodies and resting structures of other fungi such as *S. macrospora* and two different species of *Verticillium*, but not of *A. nidulans* itself (Fig. 7). This might be explained by the fact that *A. nidulans* is able to convert emodin to other structures, whereas *S. macrospora* and *Verticillium* spp. do not possess enzymes for the conversion. This kind of detoxification is one common self-protection mechanism of fungi (Keller et al., 2015). The fact that emodin represses the formation of fruiting and resting structures in such divergent fungi as *S. macrospora* and *Verticillium* spp. suggests a commonly present emodin target. Emodin is widely identified in plant families but also in fungi, particularly in *Penicillium* spp. and *Aspergillus* spp. It exhibits a wide spectrum of ecological functions and protects higher plants against herbivores, pathogens, competitors as well as extrinsic abiotic factors (Izhaki et al., 2002). In addition, emodin has pharmaceutical properties, such as anti-inflammatory and anti-tumor activities in mammalian cells by affecting the cellular NF-κB and MAPK (mitogen-activated protein kinase) signaling pathways (Huang et al., 2004; Wang et al., 2006; Xie et al., 2019). The MAPK signaling pathways are conserved from yeast to human playing a central role in transducing extracellular stimuli into intracellular responses (R. E. Chen et al., 2007; Meskiene et al., 2000). It is well known that MAPK pathways are commonly involved in fungal development (Ö. Bayram et al., 2012a). Therefore, the fungal MAPK signaling pathways might be suitable emodin targets, disordering the signaling transduction and inhibiting the sexual fruiting body or resting structure formation. We suggest that the *mdp* metabolite emodin is produced to compete other fungi in order to maintain enough space and nutrients for the own sexual fruiting body development.

Besides emodin, other *mdp*/*xpt* SMs seem to be involved in the repression of fruiting and resting structure formation, since SMs of Δ*mdpD*, which do not contain emodin, are involved in repression in *S. macrospora, V. dahliae* and *V. longisporum* (Fig. 6). Further, enriched intermediates in the *mdp*/*xpt* deletion strains Δ*mdpC*, Δ*mdpL*, Δ*mdpD*, Δ*xptA* and Δ*xptB* delayed the fruiting body maturation of *A. nidulans* (Fig. 5a). Whereas the delay was gradually rescued along with the decrease of accumulated intermediates in Δ*mdpL*, Δ*mdpD*, Δ*xptA* and Δ*xptB* (Fig 2B and Fig. S5), Δ*mdpC* showed an additional reduction in cleistothecia size even after 25 days (Fig. S7). Emodin and ω-hydroxyemodin do not show an effect on cleistothecia size when externally added to *A. nidulans* wildtype (Fig. 7). Therefore, 2,ω-dihydroxyemodin or a combination of the different anthraquinone SMs might be the responsible compounds for the reduction in cleistothecia size. Smaller cleistothecia have lower energy and material costs. The accumulation of those intermediates might be an internal signal for incomplete xanthone/arugosin biosynthesis, which leads to fruiting bodies unprotected from animal predators.

In conclusion, our results suggest that the *mdp*/*xpt* pathway in *A. nidulans* was recruited for the protection of fruiting bodies from predation. This occurred by (i) connecting the anthraquinone producing *mdp* cluster with *xpt* genes converting anthraquinones into xanthones with antifeedant activity; (ii) expressing the *mdp*/*xpt* pathway in Hülle cells that surround the fruiting bodies; and (iii) coordinating *mdp*/*xpt* expression with sexual development by the velvet complex. These findings support a new role of Hülle cells as protective cells. They establish a secure niche for *A. nidulans* by accumulating metabolites with antifeedant activity that protect the reproductive and overwintering structures from animal predators (Fig. 9), which guarantees a long-term survival of *A. nidulans*.

**Fig 9.**
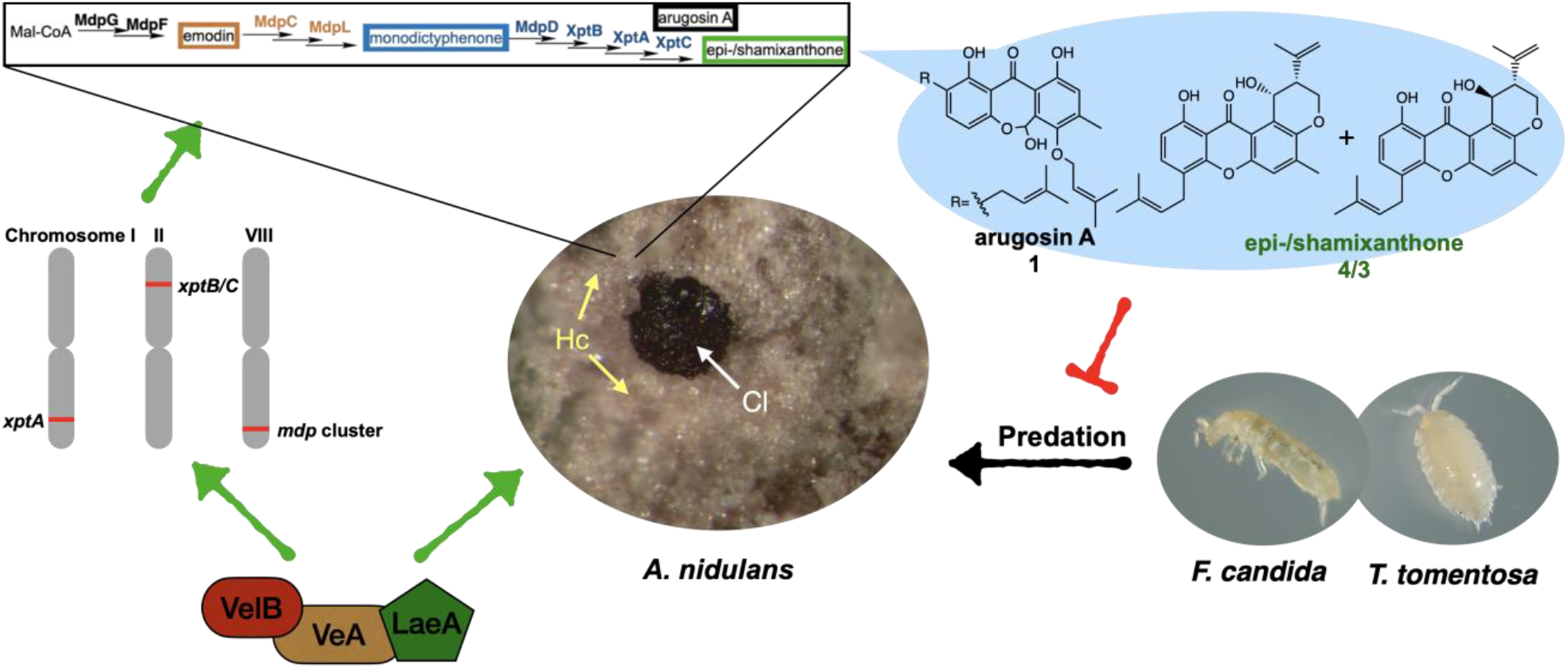
The *mdp*/*xpt* cluster metabolites establish a secure niche for *A. nidulans*. The velvet complex VelB-VeA-LaeA regulates sexual development of *A. nidulans* and the expression of *mdp/xpt* genes. The resulting metabolites are accumulated in the sexual fruiting body (Cl) nursing Hülle cells (Hc). The final products arugosin A **1** and epi-/shamixanthone **4**/**3** protect sexual fruiting bodies from the animal predators *F. candida* and *T. tomentosa*.

## Materials and Methods

### Strains and culture conditions

The *Escherichia coli* strain DH5α (Woodcock et al., 1989) was used for expression of recombinant plasmids in this study. The culture medium was <u>L</u>ysogeny <u>B</u>roth medium (LB) (Bertani et al., 1951), containing 1% bacto tryptone, 1% NaCl and 0.5% yeast extract, pH 7.5, at 37°C. The solid LB medium was added 2% agar additionally. 100 µg/ml of ampicillin was added as selective agent. *A. nidulans* strains were cultivated in <u>m</u>inimal <u>m</u>edium (MM) (AspA (70 mM NaNO_3_, 11.2 mM KH_2_PO_4_, 7 mM KCl, pH 5.5), 1% (w/v) glucose, 0.1% (v/v) Hutner’s trace elements) (Käfer et al., 1977). 2% agar was added for the solid MM plates. For selection of *A. nidulans* transformants, phleomycin (80 µg/ml) was added. *A. nidulans* FGSC A4 was used for proteome analysis. *A. nidulans* AGB552 (Ö. Bayram et al., 2012a) was used for the construction of *A. nidulans* mutants. Cultivation of strains based on AGB552 required the addition of 0.0001% 4-aminobenzoic acid.

For vegetative growth, *A. nidulans* spores were inoculated and shaken 20 h in liquid MM at 37°C in flasks with baffle. For asexual development, *A. nidulans* spores were incubated on solid MM plates at 37°C under illumination. For sexual development, the solid MM plates were sealed with Parafilm^®^ M (Merck, Darmstadt, Germany,) at 37°C in dark. Conidia were stored in 0.96% NaCl buffer with 0.02% Tween-80 (Sigma-Aldrich Chemie GmbH, Taufkirchen, Germany) at 4°C.

*Sordaria macrospora* was cultivated on cornmeal malt fructification medium (BMM) agar plates at 27°C (Dirschnabel et al., 2014). *Verticillium* spp. were cultivated on simulated xylem medium (SXM) agar plates at 25°C (Hollensteiner et al., 2017). All fungal strains are listed in table S3.

### Plasmid and *Aspergillus nidulans* strain construction

The GeneArt^®^ Seamless Cloning and Assembly Enzyme Mix (Thermo Fisher Scientific, MA, USA) or the GeneArt^®^ Seamless Cloning and Assembly Kit (Thermo Fisher Scientific) was used for the assembly of DNA fragments with the backbone. For DNA amplification, gDNA from *A. nidulans* AGB552 strain was used. The constructed and used plasmids in this study are present in table S4. The used primers are present in table S5.

The recyclable marker cassette-containing plasmid pME4319 was used as the cloning vector, which harbors the *bleo* gene conferring resistance to phleomycin (Drocourt et al., 1990). For pME4319 construction, the pBluescript II (+) vector was amplified with primers flip1, carrying a *Swa*I site, and flip2, carrying a *Pml*I site. The recyclable phleomycin resistance marker was cut from pME4305 with *Sfi*I (Thieme et al., 2018). Both fragments were assembled as described above. For the *Aspergillus* deletion strain generation, approximately 1.5 kb of the 5’ flanking region of the gene of interest, carrying a *Pme*I site, was inserted into the *Swa*I site of pME4319 and approximately 1.5 kb of the 3’ flanking region of the gene of interest, carrying a *Pme*I site, was inserted into the *Pml*I site of pME4319. For ‘on-locus’ transformation, the cassette containing the 5’ flanking region, the recyclable phleomycin resistance marker and the 3’ flanking region was cut as a linear fragment by *Pme*I.

5’ flanking region (1.5 kb, primers LL139/140) and 3’ flanking region of *mdpG* (1.6 kb, primers LL141/142) were inserted into the *Swa*I and *Pml*I sites of pME4319, respectively, resulting in pME4842. The cassette of Δ*mdpG* was excised and integrated into AGB552 for the *mdpG* deletion strain AGB1236. 5’ flanking region (1.0 kb, primers LL197/198) and 3’ flanking region of *mdpF* (1.0 kb, primers LL199/200) were integrated into pME4319, resulting in pME4843. The cassette of Δ*mdpF* was integrated into AGB552 for the *mdpF* deletion strain AGB1237. 5’ flanking region (1.0 kb, primers LL223/224) and 3’ flanking region of *mdpC* (1.1 kb, primers LL225/226) were integrated into pME4319, resulting in pME4844. The deletion cassette was integrated into AGB552 for the *mdpC* deletion strain AGB1238. 5’ flanking region (1.2 kb, primers LL193/194) and 3’ flanking region of *mdpL* (1.2 kb, primers LL195/196) were inserted into pME4319, giving rise to pME4845. The Δ*mdpL* cassette was integrated into AGB552 for the *mdpL* deletion strain AGB1239. 5’ flanking region (1.1 kb, primers LL162/163) and 3’ flanking region of *mdpD* (1.3 kb, primers LL164/165) were cloned into pME4319, giving rise to pME4846. The deletion cassette was integrated into AGB552 for the *mdpD* deletion strain AGB1240. 5’ flanking region (1.3 kb, primers LL211/212) and 3’ flanking region of *xptA* (1.3 kb, primers LL213/214) were cloned into pME4319, resulting in pME4847. The Δ*xptA* cassette was integrated into AGB552 for the *xptA* deletion strain AGB1241. 5’ flanking region (1.5 kb, primers LL215/216) and 3’ flanking region of *xptB* (1.3 kb, primers LL217/218) were cloned into pME4319, resulting in pME4848. The Δ*xptB* cassette was excised and integrated into AGB552 for the *xptB* deletion strain AGB1242. 5’ flanking region (1.0 kb, primers LL219/220) and 3’ flanking region of *xptC* (1.1 kb, primers LL221/222) were integrated into pME4319, giving rise to pME4849. The cassette of Δ*xptC* was excised and integrated into AGB552 for the *xptC* deletion strain AGB1243.

For the construction of the *mdpG* complementation strain, 1.5 kb of the 5’ flanking region as well as the *mdpG* gene were amplified in two PCRs with primers LL59/60 and primers LL61/62. Both fragments were inserted into the *Swa*I site of pME4319. 1.6 kb of 3’ flanking region of *mdpG* (primers JG1076/EFS46) was inserted into the *Pml*I site, resulting in pME4850. The *mdpG* complementation cassette was excised from pME4850 and integrated into AGB1236 for AGB1248. For construction of *mdpC* complementation strain, 1.9 kb of 5’ flanking region and the *mdpC* ORF (primers LL223/236) was inserted into the *Swa*I site of pME4319. 1.1 kb of 3’ flanking region of *mdpC* (primers LL225/226) was inserted into the *Pml*I site, giving rise to pME4851. The complementation cassette of *mdpC* was integrated into AGB1238 for AGB1249.

For the construction of *xptC:gfp*, 3.5 kb of 5’ flanking region and the *xptC* ORF was amplified with the primers BD113/114 and fused with *gfp* (primers BD106/107). The fragment was inserted into the *Swa*I site of pME4319. 3.8 kb of 3’ flanking region amplified with the primers BD115/116 was inserted into the *Pml*I site, giving rise to pME4645. The *xptC:gfp* cassette was introduced into AGB552, resulting in AGB1088.

For the construction of AGB1310, the *veA* deletion cassette was excised from pME4574 with *Pme*I and integrated into AGB552. For the construction of AGB1311, the *velB* deletion cassette was excised from pME4605 with *Pme*I and integrated into AGB552.

For the construction of pME4636, 2.3 kb of *laeA* 5’ flanking region was amplified with primers BD45/BD46 and 2.1 kb of *laeA* 3’ flanking region was amplified with primers BD47/BD48. The fragments were inserted into pME4319. The *laeA* deletion cassette was excised with *Pme*I from pME4636 and integrated into AGB552, resulting in AGB1073.

### Transformation of *E. coli* and *A. nidulans*

*E. coli* transformation was performed using the heat-shock method (Inoue et al., 1990). *A. nidulans* transformation was performed using the polyethylene glycol-mediated protoplast fusion method (Punt et al., 1992). Successful transformation was further verified by Southern hybridization (Southern et al., 1975). The recyclable marker cassette was eliminated from the genome by culturing the fungus on 1% xylose MM plate (Hartmann et al., 2010).

### Semi-quantitative reverse-transcriptase polymerase chain reaction

Fungal tissues were harvested from sexual cultures after three days of incubation and ground with a table mill in powder. The RNA isolation was performed according to the instruction of the RNeasy^®^ Plant Miniprep Kit (Qiagen, Hilden, Germany). Approximately 0.8 µg of RNA was used for cDNA synthesis according to manufacturer’s instructions of the QuantiTect^®^ Reverse Transcription Kit (Qiagen). 1 µl of cDNA was used for semi-quantitative reverse-transcriptase polymerase chain reaction. The used primers are listed in the table S6. Measurements were conducted in three independent biological replicates.

### Protein extraction of *A. nidulans*

Asexual and sexual mycelia of *A. nidulans* wildtype (FGSC A4, *veA*+) were harvested from solid agar plate cultures after three, five and seven days. Vegetative mycelia were harvested from 20 hours of liquid culture. Mycelia samples were frozen in liquid nitrogen and ground with a table mill. Protein extraction was carried out as described (Ö. Bayram et al., 2012b). Hülle cells were collected from three, five and seven days-old cleistothecia, grown under sexual conditions, by rolling cleistothecia on an agar plate surface. Hülle cells were sonicated for cell disruption repeatedly (60% of sonication power for 60 seconds, in between centrifugation for 30 seconds at 4°C) and centrifuged for 10 min (13,000 rpm, 4°C). The supernatant containing proteins was harvested for further experiments. For each sample three biological replicates were prepared.

### Protein digestion with trypsin and LC-MS analysis

Approximately 80 μg of protein were separated by SDS-PAGE for 60 minutes at 200 V. The gel was stained (Neuhoff et al., 1988) and the lanes were excised and subjected to tryptic digestion (Shevchenko et al., 1996). Digested peptides were desalted by using C18 stage tips (Rappsilber et al., 2007). The peptides were resuspended in 20 μl sample buffer (2% (v/v) acetonitrile and 0.1% (v/v) formic acid). LC-MS was performed by using a Velos Pro™ Hybrid Ion Trap-Orbitrap mass spectrometer (MS) coupled to a Dionex Ultimate 3000 HPLC (Thermo Fisher Scientific) (Schmitt et al., 2017). Proteomics raw data were searched with SEQUEST and Mascot algorithms present in Proteome Discoverer 1.4 using the *A. nidulans* genome database (AspGD) (Cerqueira et al., 2013; Eng et al., 1994; Koenig et al., 2008). The search parameter for the algorithms were: 10 ppm of precursor ion mass tolerance; 0.6 Da of fragment ion mass tolerance; 2 maximum of missed cleavage sites; variable modification by methionine oxidation; fixed cysteine static modification by carboxyamidomethylation. Results filter settings: high peptide confidence; minimal number of two peptides per protein.

### Fluorescence microscopy of fusion proteins

Vegetative mycelia (20 h) as well as three-day-old sexual mycelia and Hülle cells were transferred to an object slide. Fluorescence microphotographs were taken with a confocal light microscope (Zeiss Axiolab-Zeiss AG) equipped with a QUAN-TEM: S12SC (Photometrics) digital camera and the software package SlideBook 6 (Intelligent Imaging Innovations GmbH, Göttingen, Germany).

### Immunoblotting

Extracted proteins of three-day-old sexual fungal tissues were separated by PAGE and transferred onto a nitrocellulose membrane (GE Healthcare, Wisconsin, USA) as described (Schinke et al., 2016). Ponceau staining was used as sample loading control. After blocking in 5% skim milk powder dissolved in TBS-T buffer, the first antibody α-GFP antibody (sc-9996, Santa Cruz Biotechnology Inc., CA, USA) was diluted 1:1000 in blocking solution and the second α-mouse antibody (G21234, Invitrogen AG) was diluted 1:2000 in blocking solution. The result was visualized on a Fusion-SL7 (Vilber Lourmat, Collégien, France) system. The experiments were carried out with three biological replicates.

### Extraction of secondary metabolites and LC-MS analysis

4 µl containing approximately 1,000 conidia were point-inoculated on MM agar plates and sexually grown for two, three, five, seven and 10 days. A 5.7 cm^2^ agar piece of the colonies was cut into small pieces and covered with 5 ml of ethyl acetate in 50 ml tube. Tubes were shaken at 200 rpm at room temperature for 30 min followed by 10 min highest level ultra-sonication in a Bandelin Sonorex™ Digital 10P ultrasonic bath (Bandelin Electronic GmbH & Co. KG, Berlin, Germany). 3 ml of ethyl acetate phase was transferred to a glass tube and evaporated.

SM sample was resuspended in 700 µl methanol and centrifuged for 10 min (13,000 g, 4°C) to remove particles. 500 µl of supernatant was transferred into the LC-MS vial. LC-MS was performed by using a Q ExactiveTM Focus orbitrap mass spectrometer coupled to a Dionex Ultimate 3000 HPLC (Thermo Fisher Scientific). 5 µl of SM sample was injected into the HPLC column (Acclaim™ 120, C^18^, 5 µm, 120 Å, 4.6 x 100 mm). The running phase was set as a linear gradient from 5-95% (v/v) acetonitrile/0.1 formic acid in 20 min, plus 10 min with 95% (v/v) acetonitrile/0.1 formic acid) with a flow rate of 0.8 ml/min at 30°C in addition. The measurements were performed in positive and negative modes with a mass range of 70-1050 m/z. FreeStyle™ 1.4 (Thermo Fisher Scientific) was used for data analysis.

### Monitoring of sexual development

Approximately 1,000 conidia of *A. nidulans* strains were point-inoculated on MM agar plates and cultivated under sexual conditions. Sexual fruiting body development of each strain was monitored at two, three, four, five, seven and 10 days. The development status of the cleistothecia at the colony center was recorded over time with photomicrographs. Matured cleistothecia were collected and counted, and their diameters were measured by a microscope with the software cellSens Dimension (Olympus Europa SE & Co. KG, Hamburg, Germany). Cleistothecia were broken in 100 µl 0.02% tween buffer for ascospore quantification (*n* = 10). This was performed with three biological and three technical replicates. Hülle cells were detached from five-days-old cleistothecia by rolling the fruiting bodies on agar surface. The diameter of Hülle cells was measured as described for the cleistothecia. Two independent experiments were carried out.

### Analysis of germination of Hülle cells

Hülle cells were collected from five days old cleistothecia of AGB552, Δ*mdpG*, Δ*mdpC* and Δ*mdpL* separately. Hülle cells were picked with a MSM System 300 micromanipulator (Singer Instruments) and placed on separate fresh MM plates (*n* = 40 (±1)). For germination experiments, plates were incubated for two days under light conditions at 37°C. The germination rate of detached Hülle cells was calculated from the visible colonies after 48 hours. This was performed in two biological replicates.

### Effect of secondary metabolite on fungi

SMs of *mdp*/*xpt* gene deletion strains and *A. nidulans* wildtype AGB552, extracted from nine point-inoculated colonies, were dissolved in 450 µl of methanol. 30 µl of mixed solution was loaded on the filter paper disc (Φ = 9 mm) individually. 30 µl of pure methanol was used as a blank control. 75 µg of pure ω-hydroxyemodin (ChemFaces, Wuhan, China), emodin (VWR, Darmstadt, Germany) and chrysophanol (VWR) were dissolved in methanol and loaded on the paper disc for following tests.

Loaded paper discs were placed on plates inoculated with spores of the tested fungi. For *A. nidulans* wildtype AGB552, 1×10^5^ fresh conidia were spread on 80 ml MM agar plates and incubated under sexual growth inducing conditions for five days. The size and amount of cleistothecia on each paper disc were monitored. For *Sordaria macrospora*, 2×10^5^ spores were spread on cornmeal malt fructification medium (BMM) agar plates and grown for seven days at 27°C. For *Verticillium* spp., 1×10^5^ spores were separated completely on SXM agar plates and grown for 10 days at 25°C.

### Animal food preference

*Tenebrio molitor* larvae were purchased from Zoo & Co. Zoo-Busch (Göttingen, Germany). *Folsomia candida* was provided by the Institute of Zoology (University of Göttingen, Germany), and was kept on the plaster Petri dishes (gypsum plaster: charcoal (9: 1)) (Xu et al., 2019). *Trichorhina tomentosa* was purchased from b.t.b.e. Insektenzucht GmbH (Schnürpflingen, Germany). *A. nidulans* spores were point-inoculated on 50 ml MM agar plates and incubated for five days of sexual growth.

For the experiment with *T. molitor*, fungal colony agar pieces (Φ = 2.7 cm) were cut and placed on two opposite sides of a Petri dish (140 mm in diameter). Animals (*n =* 10). were placed into the center area of the Petri dish. The number of animals on each side was counted over a period of 8 hours. The experiment was carried out with three biological and nine technical replicates.

For the experiment with *F. candida*, the *A. nidulans* colony agar pieces (Φ = 1.5 cm) were placed onto opposite sides of the plaster Petri dishes (92 mm in diameter). Approximately 20 animals were placed onto the center of the Petri dish and the number on each side was counted over a period of 24 hours. The experiment was performed with three biological and four technical replicates.

The food choice experiment with the isopod *T. tomentosa* was carried out as described for *F. candida* with *n =* 8 animals.

## Acknowledgements

We thank Verena Große, Nicole Scheiter, Gertrud Stahlhut and Ruth Pilot for technical assistance, Kerstin Schmitt and Miriam Kolog Gulko for preparing protein digestion solutions and proteome data discussions. We acknowledge support by the doctoral programs of Göttinger Graduiertenzentrum für Neurowissenschaften, Biophysik und Molekulare Biowissenschaften (GGNB) (University of Göttingen), the China Scholarship Council (CSC) and the European Union’s Seventh Framework Programme FP7/2007-2013 (grant agreement 607332). Funding was provided by the German Research Council to GHB (DFG grant BR 1502/19-1).

## Author Contributions

Conception and overall design were performed by LL, SP, PK, JG and GHB. Experiments were performed by LL, BD, EF-S, RH, CS, DEN and JG. Data acquisition was performed by LL, BD, OV and JG. Data analysis was performed by LL, BD, OV, RH, DEN, SP, PK, JG and GHB. The manuscript was written by LL, JG and GHB with contributions of all authors.

## Additional information

**Supplementary information** is available for this paper.

**Correspondence** and request for materials should be addressed to GHB and JG.

